# DOES EXPERIENCE MODULATE AUTOMATIC IMITATION? A NEW LOOK

**DOI:** 10.1101/2024.04.17.589868

**Authors:** Francesca Genovese, Martina Fanghella, Corrado Sinigaglia, Guido Barchiesi

**Author notes:** Corresponding author Guido Barchiesi Department of Philosophy Università degli Studi di Milano Via Festa del Perdono 7, 20122 Milano, Italy. guido.

## Abstract

Automatic imitation is a stimulus-response compatibility effect wherein observing an action automatically influences motor performance. However, the mechanism underlying this effect remains controversial. Associative Sequence Learning suggests that automatic imitation arises from contingent visual and motor activity associations. Prior studies have shown that exposure to counter-imitative training can alter these visuomotor associations, suggesting that automatic imitation can be modulated by experience. Here, we aim to bring new insight into how this modulation occurs by exploring the time course of automatic imitation before and after counter-imitative training. If automatic imitation is merely a result of contingent associations, as previously suggested, the effect should consistently be modulated following such training. However, if this is not the case, automatic imitation is not, or not only, a matter of contingent association, at least not as currently understood.

## 1. INTRODUCTION

Automatic imitation is a stimulus-response compatibility effect, wherein observing another’s action heavily affects the observer’s motor performance. In a typical automatic imitation task, participants are visually presented with one of two actions (task-irrelevant stimuli) and a cue (colors or numbers) indicating the action they have to perform (task-relevant stimuli). Participants usually produce the cue commanded action faster if the latter is similar to the observed action. Such an effect has been replicated several times since its discovery, employing different experimental variants (Brass et al., 2000, 2001; Stürmer et al., 2000; for a review, see Cracco et al., 2018 and Heyes, 2011). Automatic imitation has been explained through various mechanisms. The Ideomotor theory posits a tight relationship between perception and action (Greenwald, 1970; Prinz, 1990, 2005). It claims that representing an action’s perceptual consequences is crucial for controlling its execution, as it enables the agent to anticipate the expected sensory feedback when the action is performed correctly. The same mechanism also accounts for automatic imitation: the perception of an event similar to the events one has experienced following one’s own actions facilitates executing that action.

Similarly, the Motor Resonance theory suggests that when observing another’s action, the sensory information concerning that action is automatically transformed into the motor representation involved in planning and executing that action (Rizzolatti & Sinigaglia, 2010, 2016). This explains the automatic imitation effect, with the participants performing the observed action faster even if the sensory information concerning that action is task-irrelevant.

The Associative Sequence Learning (ASL) theory moves in the opposite direction by claiming instead that automatic imitation relies on mere sensorimotor contingencies (Catmur et al., 2009; Cook et al., 2014; Heyes, 2010). According to ASL, the links between perceptual and action events develop through bidirectional Pavlovian-like associations between sensory and motor experiences. These associations are purely contingent and arise whenever actions happen to be performed and observed together without prioritizing action execution over action observation (Cook et al., 2014). The contingent co-occurrence of similar perceptual and action experiences would explain why observing an action might enhance its execution. However, this enhancement would be reduced or disappear if the contingencies differed (Ray & Heyes, 2011).

In a demonstration that has since become classic, Heyes et al. (2005) provided the first evidence that automatic imitation can be affected by contingent sensorimotor experiences. Two groups of participants underwent two distinct visuomotor training sessions. In the “imitative” training, participants watched a hand extend or flex its fingers and mimicked the observed action. In the “counter-imitative” training, participants were instructed to perform the opposite action to what they saw, flexing their fingers when the hand extended them, and vice versa. Following the training, both groups participated in a single-choice version of an automatic imitation task, extending or flexing their fingers. The training sessions had differing effects on the magnitude of automatic imitation, consistent with ASL predictions. While the imitative group exhibited an automatic imitation effect, the counter-imitative group abolished this effect.

ASL predictions have also been tested at the neurophysiological level through a passive observation paradigm. Catmur et al. (2007) employed transcranial magnetic stimulation (TMS) to stimulate the primary motor cortex and record motor-evoked potentials (MEP) from the first dorsal interosseus (FDI) and abductor digiti minimi (ADM) muscles while participants passively observed index and little finger movements involving those muscles. Before and after imitative training, MEPs in the ADM and FDI muscles were greater when observing little-finger and index-finger movements, respectively. Following counter-imitative training, the MEP amplitude pattern reversed.

Although these findings demonstrated that contingent sensorimotor experiences could modulate automatic imitation, it is still unclear how this modulation occurs. Indeed, two TMS studies on passive action observation produced contrasting results. Barchiesi and Cattaneo (2013) delivered single TMS pulses on the primary motor cortex at four intervals from the onset of a visually presented action, showing that counter-imitative training affected visuomotor facilitation only at late time points, leaving earlier ones unaffected. This result was interpreted as the interaction between two mechanisms: a fixed visuomotor transformation responsible for the early motor facilitation effects and a flexible rule-based visuomotor association for the later effects. Conversely, Cavallo et al. (2014) found that the counter-imitative training impacted visuomotor facilitation at all time points, where it was found in the pre-training session, thus suggesting that the facilitation effect would be due to only one mechanism, which is contingent association.

The present study aims to shed new light on the automatic imitation effect and its potential modulation by describing, on the behavioral front, the high-resolution time course of this effect before and after imitative and counter-imitative training. We leveraged the time course design logic implemented in Barchiesi & Cattaneo (2013) and Cavallo et al. (2014), employing a double-choice version of the automatic imitation paradigm in which the delay between the task-irrelevant event (the observation of a hand flexing or extending its fingers) and the task-relevant event (a colored cue indicating whether the action to be performed was a flexion or an extension of the fingers) was parametrically modulated.

Our main aim was to assess, for the first time, whether counter-imitative training consistently affects the automatic imitation effect across nine intervals between the presentation of the colored cue and the observed action, as reported during a pre-training session. Suppose automatic imitation entirely depends on sensorimotor contingencies, as assumed by ASL and suggested by Heyes et al. (2005)’s findings. In that case, one should expect that the counter-imitative training causes a suppression or (at least) a reduction of the whole time course of the automatic imitation effect. If this is not the case, and the automatic imitation effect remains unaffected by counter-imitative training at any point during its progression, it would involve that the automatic imitation effect is not, or not only, a matter of contingent association, at least not in the way it has been understood thus far.

## 2. MATERIALS AND METHODS

### 2.1 Sample Size Estimation

Three previous behavioral experiments implementing a “training-test” rationale employed 10 (Heyes et al., 2005), 8 (Press et al., 2007), and 12 (Cook et al., 2010) participants for each training group. To detect even small effects resulting from our experimental design, we decided to roughly quadruplicate the number of participants for each training group to 40.

The result of a sensitivity analysis performed in GPower (alpha = 0.05, beta = 0.8, correlation among repeated measures = 0.5, non-sphericity correction = 1, number of groups = 2, number of measurements = 36 reflecting 2 trainings x 2 sessions x 9 delays) indicates that employing a total of 80 participants for our experimental design is sufficient for detecting effects as small as Cohen’s f = 0.067, which is typically considered “small” (Faul et al., 2007; Perugini et al., 2018).

### 2.2 Participants

We collected data from 84 participants (right-handed, with no neurological or psychiatric disorders, vision either corrected or normal); four of them did not participate in all the experimental sessions and were excluded from the analyses. Two groups of 40 participants were enrolled in the “Counter” experiment (17 women aged 18-50 years) or the “Imitative” experiment (17 women aged 18-30 years). The study complied with the revised Helsinki Declaration (World Medical Association General Assembly, 2008) and was approved by the local Ethical Committee. All participants gave written informed consent for their participation.

### 2.3 Stimuli

Four right-hand pictures on a black background (two male and two female hands) were presented in the experimental sessions. Each hand could be presented with the fingers in three positions: flexed, intermediate (neutrally positioned), or extended. Building on previous research (Barchiesi & Cattaneo, 2015; Heyes et al., 2005; Ubaldi et al., 2015), we leveraged the illusion of movement produced by the rapid transition from images of a neutrally positioned hand to those showing flexed or extended fingers. Two small, round objects were displayed respectively between the fingers and externally on the dorsal side of the presented hand; when a flexion of the fingers was displayed, the action was a goal-directed grasp, while when the fingers extended, the displayed action was a goal-directed touch of the external round object with the dorsal portion of the tips of the fingers.

### 2.4 Response Detection System

#### 2.4.1 Flex-sensor

During each experimental session, participants had to produce a flexion or an extension of right hand fingers following the appearance of a colored cue on the screen (see next paragraph). In order to detect such responses, we employed a 95 mm long bending variable resistor (from here on, “flex-sensor”) placed on the participant’s right index finger. To this aim, two soft plastic sheaths (∼0.5 cm diameter x ∼2.5 cm length) were attached on the dorsal side of the proximal and middle phalanxes of the right-hand index finger and kept in place with skin adhesive tape. Afterward, the flex-sensor was inserted into the sheats and then fixed onto the index metacarpus so that when participants flexed or extended their fingers, the flex-sensor bent and slid inside the sheaths. When the fingers relaxed, the flex-sensor could return to its original position (Supplementary Material Figures 1, 2, and 3).

The flex-sensor was connected in series to a fixed resistor within a voltage divider circuit so that the finger bending produced voltage modulations across the poles of the flex-sensor; to record the data, we employed an Arduino® board (which powered the whole circuit as well). We transformed the voltage output into conductivity values across the poles of the flex-sensor. In order to normalize the circuit output according to the degree of each participant’s finger flexion/extension range, the conductivity output values detected during maximal flexion and extension of the fingers were marked and mapped respectively to 0 and 1000 arbitrary units. This procedure was repeated for each experimental session. To detect participants’ responses, extension and flexion thresholds were set to be equal to the conductivity detected at the onset of the colored cue plus/minus 100 arbitrary units; for example, if the initial finger position was detected as 430 arbitrary units, 530 and 330 would have been the thresholds for extensions and flexion, respectively. During each session, the participant’s right hand was covered by an opaque box, preventing them from seeing their hand moving.

#### 2.4.2 Foam Rubber Manipulandum

Between the participants and the PC monitor, a foam rubber structure (diagonal ∼4,5 cm, height ∼14 cm) and a rigid base on the right side were fixed on the table. Participants had to lay the ulnar side of their right hand on the rigid base and place the fingers around the foam rubber structure. This way, the thickness of the foam rubber structure kept the fingers in an intermediate position between extension and flexion and at the same time its softness allowed the fingers to flex (supplementary material Figure 2).

### 2.5 General Experimental Structure

Each participant underwent three sessions: a “Pre-training” session (common to both groups), in which a forced double-choice reaction time version of the automatic imitation paradigm was tested. A training session followed the Pre-training: in this session, one group performed a counter-imitative visuomotor training (“Counter” group) while the other performed an imitative visuomotor training (“Imitative” group). Both groups eventually performed a “Post-training” session, similar to the Pre-training, the following day.

#### 2.5.1 Automatic Imitation Time Course (Pre-training and Post-training Sessions)

##### 2.5.1.1 Task and Trial Structure

The experimental design was implemented and run by custom Matlab R2021b (Mathworks®) scripts employing Psychtoolbox functions (Brainard, 1997; Kleiner et al., 2007; Pelli, 1997). Stimuli were presented on a 60 Hz refresh rate monitor (DELL P2419H, resolution 1920 x 1080 pixels, screen size: 23.8 inches). Each trial presented a randomly oriented (0-360°) neutrally positioned hand on the screen, which lasted randomly between 1000 and 1500 ms. In the remainder of the trial, a colored cue (either light green or purple, lasting 67 ms) and an action (either a flexion or an extension of the fingers, hereafter referred to here as “observed action”) were presented on the screen; the critical feature of each trial was the temporal delay between the presentation of the colored cue and the presentation of the observed action. Here we conceptually grouped them into three categories: in the “negative delays” trials, the neutrally positioned hand turned colored, and after 700, 300, 200, or 100 ms, an action was presented on the screen; in the “simultaneous delay” trials, the colored cue was presented simultaneously with the observed action (0 ms delay); eventually in the “positive delays” trials, the colored cue could be presented 100, 200, 300, 400 ms after the observed action (Figure 1).

**Figure 1:**
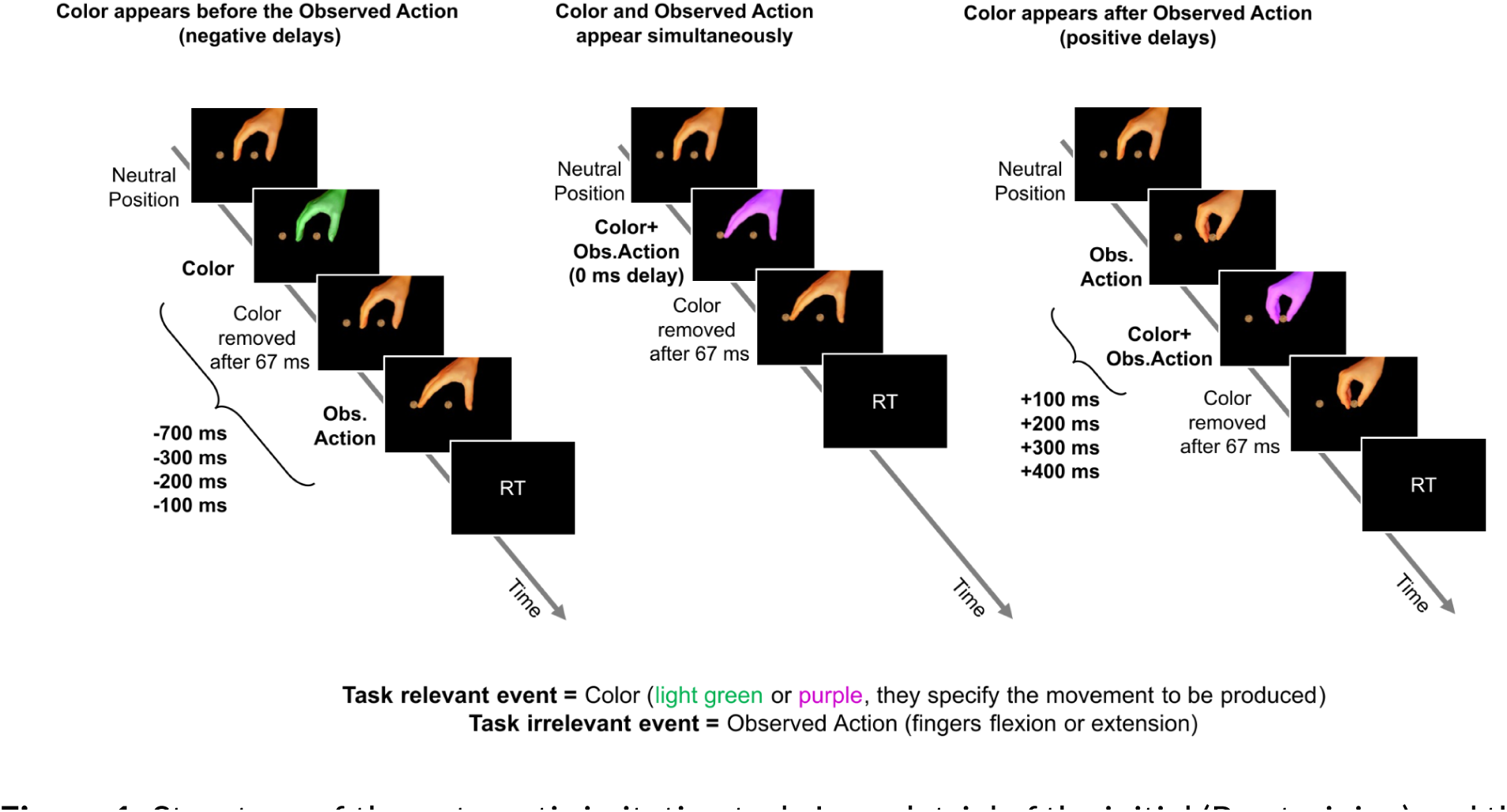
Structure of the automatic imitation task. In each trial of the initial (Pre-training) and the last session (Post-training), the task-relevant event, the colored cue (either light green or purple), appears at one of nine different delays with respect to the task-irrelevant event, the observed action (either fingers flexion or extension). The order of the delays has been randomized across trials. The leftmost timeline depicts a trial where the colored cue flashes on the neutrally positioned hand -700, -300, -200, or -100 ms relative to the subsequent presentation of the observed action (“negative delays trials”). The middle timeline represents a trial where the colored cue and the observed action appear simultaneously (delay = 0 ms). The rightmost timeline represents a trial in which the colored cue appears 100, 200, 300, or 400 ms after the observed action has been presented on the screen (“positive delays trials”). In each trial type, the colored cue lasts 67 ms. The orientation of the displayed hand was randomized across trials between 0 and 360 degrees.

The order of the nine delays was randomized across sessions and participants. The participants’ task was to produce a flexion or an extension of their right hand fingers according to the presented colored cue as fast as possible and as accurately as possible; the association rule between the color and the movement to be produced was counterbalanced across participants. After either a detected response or the end of the response time window (until 700 ms from the colored cue presentation), participants were provided with a feedback screen for 500 ms: if the response was correct and provided between 150 and 700 ms, their reaction time in milliseconds was displayed; if the detected reaction time was faster than 150 ms, a “Do not guess!” alert was displayed, if it exceeded the deadline time, a “Too Slow” alert was presented; if the provided response was wrong, a “Wrong!” label was shown (feedback labels have been displayed in Italian). Eventually, a black screen was displayed for 1000 ms before the subsequent trial started.

To enhance participants’ motivation and encourage them to perform at their highest level, we informed them, at the onset of each session, that the best performer would receive a prize at the end of data collection.

Each Pre-training and Post-training session comprised 720 trials, with 80 trials for each of the nine delays. Within each delay, there were 40 congruent trials, where the observed action matched the movement commanded by the colored cue, and 40 incongruent trials, where the observed actions were the opposite of those commanded by the colored cue. Participants could briefly rest within each session once every 240 trials (2 breaks). Since the deadline for response production was set at 700 ms from the colored cue appearance, in the -700 ms delay, the action on the screen was shown for only one frame, and the same applied for all the trials in which the responses were produced before action observation.

On each trial, at the onset of the colored cue, a TTL trigger was delivered through the parallel port of the presentation PC into a digital input pin of an Arduino board. A custom-made routine, loaded on the board, measured the value of the flex-sensor output and compared it with the flexion and extension thresholds for response detection (see Flex-sensor paragraph). Once the response and the reaction time were calculated, they were sent back, via serial communication, to Matlab for visual feedback presentation.

##### 2.5.1.2 Warm-up

Before the Pre-training session, participants underwent a brief “warm-up” session (∼ 5 minutes, 80 trials) to familiarize themselves with the association between the movements to be produced and the colored cues. The task was identical to the one employed in the Pre-training and Post-training sessions, with the difference that the hands on the screen did not move (they only changed color) and that the time window for response production was increased to 3000 ms. Participants underwent the warm-up session only before the Pre-training session.

#### 2.5.2 Counter-imitative Training and Imitative Training Sessions

The Counter and Imitative group training sessions comprised 720 trials divided into 10 consecutive blocks. The trial structure was similar to the Pre-training and Post-training sessions but with a few differences: the hands were oriented from an egocentric perspective, and the fingers were either extended or flexed without changing color. Notably, the participants’ task in the Counter group was to produce the opposite action compared to the one they observed, while in the Imitative group, it was to produce the same action as the one observed. Each participant underwent only one of the two trainings, depending on the group they were assigned to. As in the Pre-training and Post-training sessions, participants were instructed to produce their responses as quickly and accurately as possible. The feedback screens were the same as in the Pre- and Post-training sessions.

### 2.6 Data Analysis

#### 2.6.1 Analysis: Pre and Post-training Sessions

To understand whether the automatic imitation effect was differently affected by the two trainings, a mixed three-way 2×9×2 ANOVA has been conducted with SESSION (Pre-training and Post-training) and DELAY (−700, −300, 200, 100, 0, 100, 200, 300, and 400 ms) as within-subject factors and TRAINING (Counter and Imitative) as a between-subject factor. In order to facilitate the interpretation of the results, an index of automatic imitation effect was computed as the dependent variable: we calculated the median RT of the congruent trials, and we subtracted it from the median RT of the incongruent ones separately for each cell of the three-way ANOVA and for each participant. Thus, if participants were slower in the incongruent trials than congruent ones, this would have been reflected as a positive automatic imitation index, and vice versa if incongruent trials were faster than congruent ones.

Given that a significant three-way interaction was detected, we separated the three-way design into two two-way 2×9 repeated-measures ANOVAs (one for each training group), with SESSION and DELAY as within-subject factors, to understand whether the training changed the automatic imitation effect in the Post-training compared to the Pre-training session.

A significant two-way interaction was only obtained for the Counter group. Since the experiment aimed to describe how the time course of the automatic imitation effect was modulated between the Pre- and Post-training sessions, we first identified, within the Pre-training session, at which delays an automatic imitation effect was detected; secondly, we tested if, at these delays, the automatic imitation effect was modulated in the Post-training session.

Thus, we performed a one-way repeated-measures ANOVA on the Counter group’s Pre-training session with DELAY as the only factor, followed by nine Bonferroni-corrected t-tests (0.05/9) at each delay.

Finally, using five Bonferroni-corrected (0.05/5 = 0.01) paired sample t-tests, we tested the modulation of the automatic imitation effect between the Post- and the Pre-training session at those delays where an automatic imitation effect was detected in the Pre-training session.

As a post-hoc analysis, we tested a residual automatic imitation effect within the Post-training session using nine Bonferroni-corrected post-hoc tests across delays.

Given the absence of a two-way interaction within the Imitative group, we did not further analyze the modulation of the automatic imitation effect between the Pre- and Post-training sessions. However, for the sake of Figure 2 clarity and interpretability, we performed nine Bonferroni-corrected paired-sample t-tests for each session across all delays also for the Imitative group (see Figure 2).

**Figure 2:**
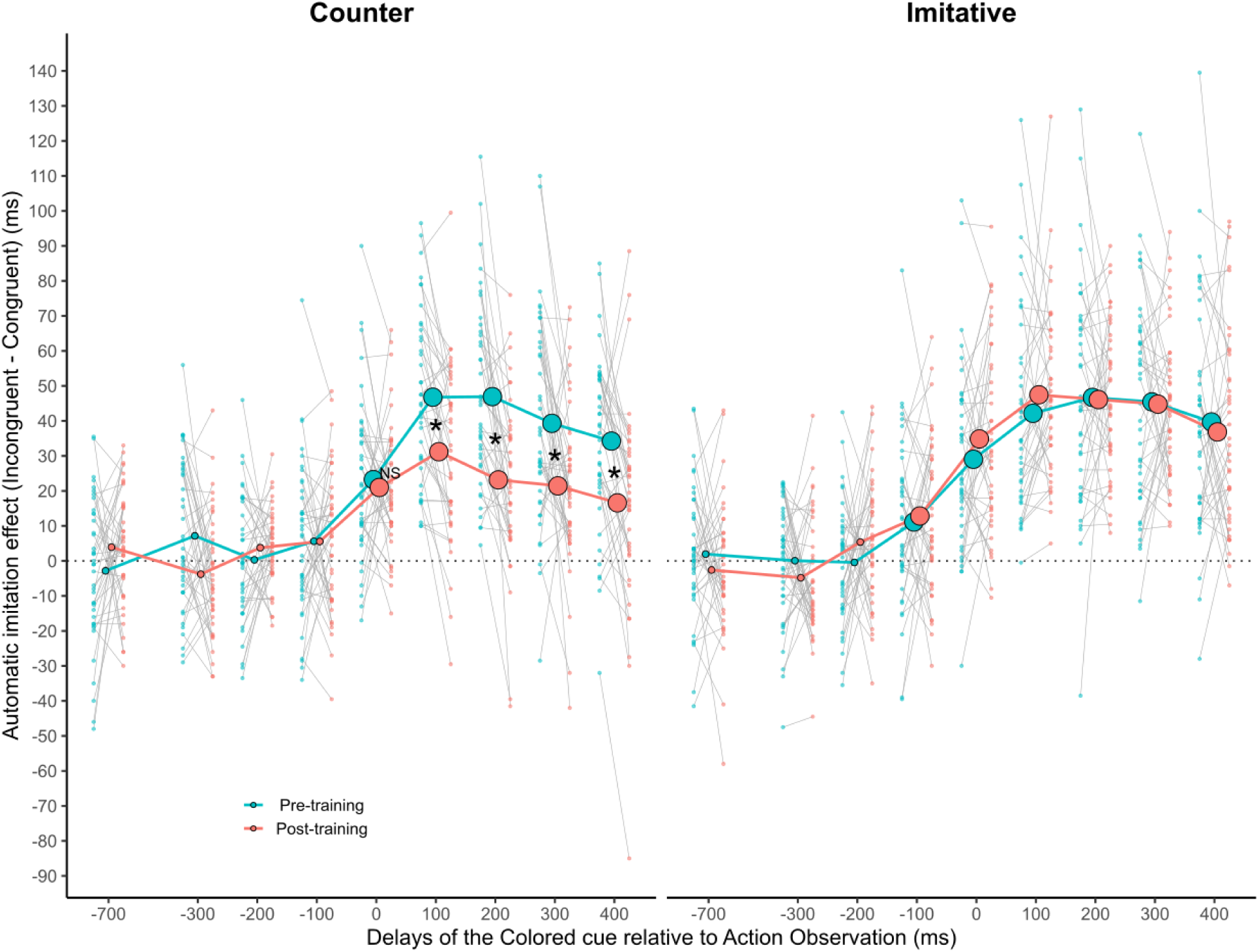
The plot shows the average time course of the automatic imitation effect (RTs from incongruent trials minus RTs from congruent trials) within the Counter group (left) and the Imitative group (right), both in the Pre-training (thick turquoise line) and Post-training session (thick salmon line). Dots on thick lines represent delays at which the automatic imitation effect is significantly greater than zero (Bonferroni corrected for nine comparisons separately for each session and each group). For each delay where a significant automatic imitation was found in the Pre-training session, we compared the magnitude of the automatic imitation effect between Pre-training and Post-training sessions. The NS label signals a non-significant difference between Pre- and Post-training sessions. In contrast, the “*” symbol signals a significant difference in automatic imitation magnitude between the two sessions (Bonferroni corrected for five comparisons). No labels are present in the Imitative group plot since no two-way interaction has been detected; indeed, no statistical testing has been conducted relating the differences between automatic imitation effects between the Pre-training and the Post-training sessions. Thin grey lines at each delay show single-subject data. Within each delay, the left side of each thin line (small turquoise dots) identifies Pre-training data, and the right (small salmon dots) identifies Post-training data.

**Figure 3:**
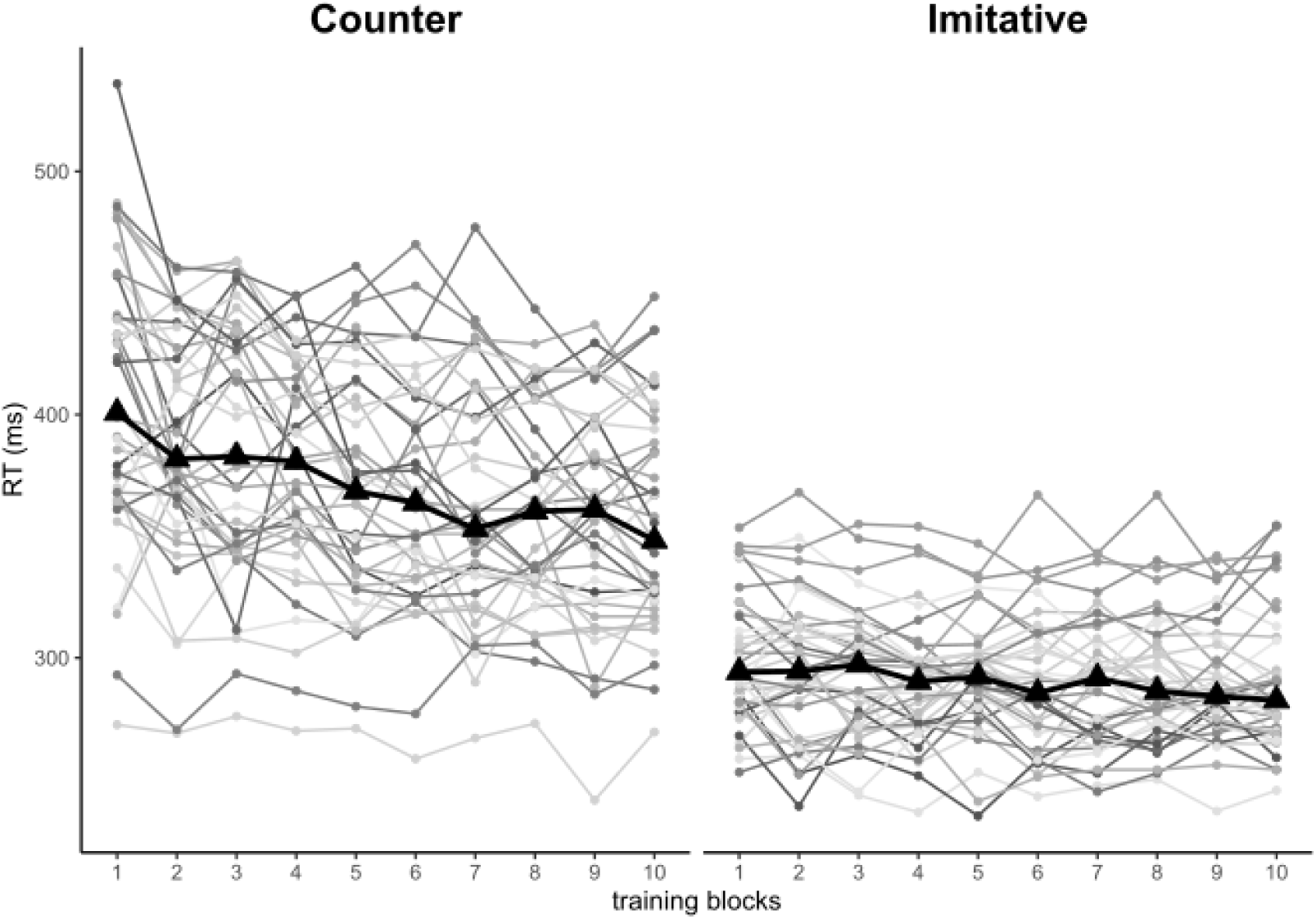
The plot represents the modulation of the RTs across the training block for the Counter group (left) and the Imitative group(right). The black triangles connected by the thick black line show the average effect of the training across the ten blocks. The grey dots connected by the thin lines of different shades of grey represent the median RT value of each participant at each training block.

#### 2.6.2 Analysis: Counter and Imitative Training Sessions

RTs between 150 ms and 700 ms from correct responses were included in the analysis. Separately for each group, we conducted a one-way repeated-measures ANOVA with TRAINING BLOCKS (from the first to the tenth) as the independent variable and the median RT for each block as the dependent variable. We fitted a linear-mixed regression model to test a linear trend across the blocks. The model predicted the RTs based on training blocks as a fixed effect, accounting for random intercepts and slopes for training blocks across subjects.

## 3. RESULTS

### 3.1 Results: Pre and Post-training Sessions

The mixed three-way ANOVA (TRAINING x SESSION x DELAY) produced a significant three-way interaction (F_6.99 545.01_ = 3.132; *p* = 0.003; partial-η^2^ = 0.039, Greenhouse-Geisser correction was applied for departure from sphericity), indicating that the automatic imitation effect in the Post-training session was affected differently by the two type of training across delays, compared to the Pre-training session (Supplementary Material Table S1).

The analysis of the two two-way repeated-measures ANOVAs (SESSION x DELAY) for each training group produced a two-way interaction for the Counter group (F_6.17, 240.76_ = 6.55; *p* < 0.001; partial-η^2^ = 0.14, Greenhouse-Geisser corrected), indicating that the automatic imitation effect in the Post-training session was differently modulated across delays with respect to the Pre-training session. We did not find such an interaction for the Imitative group (F_6.59, 256.99_ = 0.921; *p* = 0.487; partial-η^2^ = 0.023, Greenhouse-Geisser corrected), indicating that the automatic imitation effect in the Post-training session was not modulated across delays with respect to the Pre-training session (Supplementary Material Table S2).

The one-way repeated-measures ANOVA conducted on the delays of the Pre-training session of the Counter group produced a significant effect (Supplementary Material Table S3) (F_5.88, 229.31_ = 32.8, *p* < 0.001; partial-η^2^ = 0.457 Greenhouse-Geisser corrected), indicating that the automatic imitation effect was modulated across delays.

Nine Bonferroni-corrected t-tests against zero (0.05/9 = 0.0056, Table 1) detected a positive automatic imitation effect from 0 to 400 ms delays.

**Table 1.**

| Counter Group |  |  |  |  |  |  |  |  |  |  |  |  |  |  |  |
| --- | --- | --- | --- | --- | --- | --- | --- | --- | --- | --- | --- | --- | --- | --- | --- |
| Pre-training |  |  |  |  |  |  |  | Post-training |  |  |  |  |  |  |  |
| Delay (ms) | Incongruent (ms) | Congruent (ms) | Automatic Imitation (ms) | T-stat | df | Cohen's d | P-value ($p<0.0056$) | Delay (ms) | Incongruent (ms) | Congruent (ms) | Automatic Imitation (ms) | T-stat | df | Cohen's d | P-value ($p<0.0056$) |
| -700 | 384 | 387 | -3 | -0.86 | 39 | -0.14 | 0.393 | -700 | 370 | 366 | 4 | 1.50 | 39 | 0.24 | 0.141 |
| -300 | 391 | 384 | 7 | 2.12 | 39 | 0.34 | 0.040 | -300 | 366 | 370 | -4 | -1.39 | 39 | -0.22 | 0.173 |
| -200 | 383 | 383 | 0 | 0.11 | 39 | 0.02 | 0.916 | -200 | 370 | 366 | 4 | 2.19 | 39 | 0.35 | 0.034 |
| -100 | 389 | 383 | 6 | 1.61 | 39 | 0.25 | 0.116 | -100 | 370 | 365 | 5 | 1.73 | 39 | 0.27 | 0.091 |
| 0 | 396 | 372 | 24 | 6.37 | 39 | 1.01 | <.001 | 0 | 373 | 352 | 21 | 7.14 | 39 | 1.13 | <.001 |
| 100 | 394 | 347 | 47 | 12.15 | 39 | 1.92 | <.001 | 100 | 366 | 335 | 31 | 7.91 | 39 | 1.25 | <.001 |
| 200 | 383 | 336 | 47 | 11.18 | 39 | 1.77 | <.001 | 200 | 352 | 329 | 23 | 0.25 | 39 | 0.95 | <.001 |
| 300 | 375 | 336 | 39 | 8.40 | 39 | 1.33 | <.001 | 300 | 341 | 320 | 21 | 5.71 | 39 | 0.90 | <.001 |
| 400 | 367 | 332 | 35 | 8.65 | 39 | 1.37 | <.001 | 400 | 339 | 322 | 17 | 3.52 | 39 | 0.56 | 0.001 |
| Imitative Group |  |  |  |  |  |  |  |  |  |  |  |  |  |  |  |
| Pre-training |  |  |  |  |  |  |  | Post-training |  |  |  |  |  |  |  |
| Delay (ms) | Incongruent (ms) | Congruent (ms) | Automatic Imitation (ms) | T-stat | df | Cohen's d | P-value ($p<0.0056$) | Delay (ms) | Incongruent (ms) | Congruent (ms) | Automatic Imitation (ms) | T-stat | df | Cohen's d | P-value ($p<0.0056$) |
| -700 | 385 | 383 | 2 | 0.66 | 39 | 0.10 | 0.513 | -700 | 368 | 371 | -3 | -0.85 | 39 | -0.14 | 0.398 |
| -300 | 385 | 385 | 0 | 0.02 | 39 | 0.00 | 0.985 | -300 | 368 | 373 | -5 | -1.79 | 39 | -0.28 | 0.082 |
| -200 | 382 | 382 | 0 | -0.16 | 39 | -0.03 | 0.871 | -200 | 372 | 366 | 6 | 1.89 | 39 | 0.30 | 0.066 |
| -100 | 389 | 378 | 11 | 2.95 | 39 | 0.47 | 0.005 | -100 | 375 | 362 | 13 | 3.66 | 39 | 0.58 | <.001 |
| 0 | 398 | 369 | 29 | 6.90 | 39 | 1.09 | <.001 | 0 | 387 | 352 | 35 | 8.24 | 39 | 1.30 | <.001 |
| 100 | 386 | 343 | 43 | 9.11 | 39 | 1.44 | <.001 | 100 | 374 | 326 | 48 | 12.34 | 39 | 1.95 | <.001 |
| 200 | 380 | 333 | 47 | 9.13 | 39 | 1.44 | <.001 | 200 | 364 | 318 | 46 | 13.31 | 39 | 2.10 | <.001 |
| 300 | 372 | 326 | 46 | 9.50 | 39 | 1.50 | <.001 | 300 | 357 | 312 | 45 | 13.25 | 39 | 2.10 | <.001 |
| 400 | 366 | 326 | 40 | 7.60 | 39 | 1.20 | <.001 | 400 | 349 | 312 | 37 | 8.20 | 39 | 1.30 | <.001 |

We then tested whether the automatic imitation effect was reduced between the Post-training and Pre-training sessions within the Counter group. We limited the testing to those delays in which the automatic imitation was found in the Pre-training session (0, +100, +200, +300, +400 ms). The paired sample t-tests (Bonferroni corrected for five comparisons) showed that, compared to the Pre-training session, Post-training automatic imitation effects were reduced from +100 ms to 400 ms delays (all ps <0.01; see Table 2 for full results). In contrast, they were not modulated at the 0 ms delay, i.e., at the first delay in which automatic imitation emerged in the Pre-training session (t_39_= 0.52 *p* = 0.604, Cohen’s d = 0.083).

**Table 2.**

| Pre vs. Post - Automatic Imitation Effect (Counter Group) |  |  |  |  |  |  |
| --- | --- | --- | --- | --- | --- | --- |
| Delay (ms) | Pre (ms) | Post (ms) | T-statistic | df | Cohen's d | P-value<br>($p < 0.01$) |
| 0 | 23.238 | 20.987 | 0.5236 | 39 | 0.08279 | 0.604 |
| 100 | 46.775 | 31.188 | 4.4742 | 39 | 0.70744 | < .001 |
| 200 | 46.938 | 23.163 | 4.877 | 39 | 0.77112 | < .001 |
| 300 | 39.312 | 21.45 | 3.0773 | 39 | 0.48656 | 0.004 |
| 400 | 34.237 | 16.587 | 3.7388 | 39 | 0.59115 | < .001 |

Eventually, as a post-hoc analysis, we tested the (residual) presence of the automatic imitation effect in the Counter group’s Post-training session. As in the Pre-training session, the automatic imitation effect was detected from the 0 ms delay onward (Bonferroni corrected for nine comparisons) (Table 1).

We did not analyze the Imitative group’s interactions further, given the absence of a two-way interaction within the two-way ANOVA. As stated in the Analysis paragraph, we performed nine Bonferroni-corrected t-tests separately for each session across all delays also for the Imitative group. Results are displayed in Figure 2 and Table 1.

### 3.2 Results: Counter and Imitative Training Sessions

The one-way repeated-measures ANOVA conducted on the training session of the Counter group produced a main effect of the TRAINING BLOCKS variable (F_3.89,151.82_ = 19.26, *p*<0.001, partial-η^2^ = 0.33, Greenhouse-Geisser corrected). A significant linear trend has been detected across the counter-imitative training blocks, showing an average RT decrease of 5 ms per training block (estimated -5.0037, SE = 0.72, t_39_ = -6.977 p < 0.001). Also, for the Imitative group, the one-way repeated-measures ANOVA produced a main effect of the TRAINING BLOCKS variable (F_5.84, 227.85_ = 3.50, *p* < 0.003, partial-η^2^ = 0.08, Greenhouse-Geisser corrected). A significant linear trend has been detected across imitative training blocks, showing an average RT decrease of 0.97 ms per training block (estimated -0.9729, SE = 0.3137, t_38.99_ = -3.102, *p* < 0.004).

## 4. DISCUSSION

In the present study, we tested the time course of the automatic imitation effect by employing a forced-choice reaction time paradigm in which we parametrically varied the delay between the task-relevant event (a colored cue) and the task-irrelevant event (the observed action) before and after an imitative or a counter-imitative training. This allowed us to investigate whether counter-imitative training abolished or at least reduced the automatic imitation effect. In particular, we aimed to assess whether the putative abolishment or reduction was detected at every delay between the observed action and the colored cue, where automatic imitation was present during the Pre-training session.

Our results showed that the automatic imitation effect is already present when the delay between the colored cue and the action observed is 0 ms and lasts until 400 ms. Strikingly, our results highlighted that the automatic imitation effect was modulated but not abolished after counter-imitative training. We detected a consistent reduction of the automatic imitation effect in the Counter group during the Post-training session. However, this reduction occurred only when the colored cue was presented between 100 and 400 ms after the action observed. No reduction was detected at 0 ms. For the Imitative group, we did not find any modulatory effect of the imitative training between the Pre- and Post-training sessions. This suggested that the modulation caused by counter-imitative training was training-specific.

These results contrast with those of Heyes et al. (2005), who found a non-significant difference between incompatible and compatible actions after counter-imitative training. In order to fully evaluate this contrast, it is essential to remember that our and Heyes et al.’s studies differ in some critical features. We employed a test-retest strategy within each training group, while they tested only the equivalent of our Post-training session in the two groups, thus being unable to remove the between-subject variance in their design. Another important difference is that Heyes and colleagues employed a single-choice reaction time paradigm whereby the observed action was used as a go-signal, thus representing a relevant event, while in our procedure it represented the irrelevant event; this potential difference in attention allocation might have magnified the motor bias determined by the action observation (Chong et al., 2009). However, we did find an automatic imitation effect in both sessions, which was reduced at later delays of the Post-training one of the Counter group, thus suggesting that the attentional allocation was sufficient to produce both the automatic imitation effect and the reduction of the latter due to the counter-imitative training. Finally, our sample was four times greater than the one they employed, potentially making it more sensitive to slight differences between conditions.

Automatic imitation is a behavioral effect that is supposed to arise due to visuomotor mappings that facilitate the corresponding motor representation connected to the action observed (Brass et al., 2000, 2001; Stürmer et al., 2000). ASL states that these mappings arise from contingent visual and motor activity associations. When contingencies vary, the same associative mechanism producing automatic imitation in the Pre-training session abolishes or reduces this effect in the Post-training session (Catmur et al., 2009; Cook et al., 2014; Heyes, 2010).

If this were the case, we would have expected to find that, for each delay at which the automatic imitation effect was found in the Pre-training session of the Counter group, a corresponding abolishment or reduction would have also been detected in the Post-training session. However, our results indicate a reduction in the automatic imitation effect is present after a counter-imitative training session, but only at some delays where automatic imitation was found during the Pre-training session.

A dual-route model might explain the differential impact of counter-imitative training on the automatic imitation effect. Dual-route models have been applied to many different domains, ranging from attention (Schneider & Shiffrin, 1977) to decision-making (Kahneman & Frederick, 2002) and moral cognition (Greene, 2007). Although these models are often characterized in various (and sometimes incompatible) ways (Evans & Stanovich, 2013), they usually share the minimal assumption that the condition that influences whether one process or mechanism occurs diverges from the conditions that influence whether another occurs.

According to the dual-route model, the time course of automatic imitation could result from the interaction of two different visuomotor mechanisms: a rigid and faster visuomotor mapping, similar to that postulated by the Ideomotor and Motor resonance theories, that would cause the automatic imitation effect, and a more flexible and slower “rule-based” mapping (Barchiesi & Cattaneo, 2013), that the counter-imitative training session would have induced. According to this model, action observation after the counter-imitative training would trigger both the automatic imitation route, which facilitates motor responses similar to those observed, and the rule-based route, which facilitates the opposed motor responses. While the automatic imitation route is fast, the rule-based route takes more time to implement.

Similarly, Tagliabue et al. (2000) invoked a dual-route model to account for the disruption of the Simon effect caused by a previous counter-spatial compatibility session. They postulated the interaction between “long-term” and “short-term” memory links. The counter-spatial training did not modify the long-term links, as their effects resulted in the short-term links. The former conceptually corresponds to the rigid visuomotor mapping in our model, while the latter corresponds to our “rule-based” mapping.

Although the dual-route model fits our results, a contingency-based explanation for the automatic imitation effect is possible. Indeed, we cannot rule out the possibility that a (significantly) more extended training might also affect the visuomotor mechanism responsible for the early automatic imitation effect. However, the reasons why this should happen are not provided by ASL theory. Furthermore, consider the hypothesis that lifelong Pavlovian-like visuomotor contingencies are responsible for the automatic imitation effect. In that case, these visuomotor links should be expected to resist a relatively brief counter-imitative training. Contrary to what was hypothesized by ASL theory, our results suggest that the brief counter-imitative training employed in our experimental design does not modify such links. Regardless of which mechanism is responsible for the visuomotor mapping involved in the automatic imitation effect, our results showed that brief counter-imitative training is insufficient to abolish this effect, at least according to our results.

Automatic imitation exhibits a specific time course. Exploring this time course allows us to better understand whether and to what extent experience can modulate the automatic imitation effect. We provided evidence that such an effect cannot be reduced to mere sensorimotor contingencies, at least not in the way it has been understood thus far.

