## Supplementary Figures and Tables for "DOES EXPERIENCE MODULATE AUTOMATIC IMITATION? A NEW LOOK"

### Supplementary Materials

#### Table S1

| Three-way Mixed Anova | | | | | | |
|---|---|---|---|---|---|---|
| | **Sum of Squares** | **df** | **Mean Square** | **F** | ***p*** | **η²p** |
| **Session** | 5887 | 1 | 5887 | 10.05 | **0.002** | 0.114 |
| **Session ✻ training** | 7666 | 1 | 7666 | 13.08 | **< .001** | 0.144 |
| Residual | 45702 | 78 | 586 | | | |
| **Delay** | 424390 | 6 | 70788 | 94.84 | **< .001** | 0.549 |
| **Delay ✻ training** | 13833 | 6 | 2307 | 3.09 | 0.006 | 0.038 |
| Residual | 349023 | 467.63 | 746 | | | |
| **Session ✻ Delay** | 12319 | 6.99 | 1763 | 4.02 | **< .001** | 0.049 |
| **Session ✻ Delay ✻ training** | 9593 | 6.99 | 1373 | 3.13 | **0.003** | 0.039 |
| Residual | 238901 | 545.01 | 438 | | | |

#### Table S2

| Two-way ANOVAs | | | | | | | | | | | | | |
|---|---|---|---|---|---|---|---|---|---|---|---|---|---|
| Counter Group | | | | | | | Imitative Group | | | | | | |
| | **Sum of squares** | **df** | **Mean square** | **F** | ***p*** | **η²p** | | **Sum of squares** | **df** | **Mean square** | **F** | ***p*** | **η²p** |
| **Session** | 13494 | 1 | 13494 | 22.29 | **< .001** | 0.364 | **Session** | 58.7 | 1 | 58.7 | 0.104 | 0.749 | 0.003 |
| Residual | 23611 | 39 | 605 | | | | Residual | 22090.7 | 39 | 566.4 | | | |
| **Delay** | 154813 | 5.71 | 19352 | 36.02 | **< .001** | 0.48 | **Delay** | 283410 | 5.26 | 53885 | 60.936 | **< .001** | 0.61 |
| Residual | 167637 | 222.57 | 537 | | | | Residual | 181386.3 | 205.12 | 884.3 | | | |
| **Session ✻ Delay** | 18931 | 6.17 | 2366 | 6.55 | **< .001** | 0.144 | **Session ✻ Delay** | 2980.6 | 6.59 | 452.3 | 0.921 | 0.487 | 0.023 |
| Residual | 112697 | 240.76 | 361 | | | | Residual | 126204 | 256.99 | 491.1 | | | |

#### Table S3

| One-way ANOVA | | | | | | |
|---|---|---|---|---|---|---|
| | **Sum of Squares** | **df** | **Mean Square** | **F** | **p** | **η²p** |
| **Delay** | 130427 | 5.88 | 22183 | 32.8 | **< .001** | 0.457 |
| Residual | 155240 | 229.31 | 677 | | | |

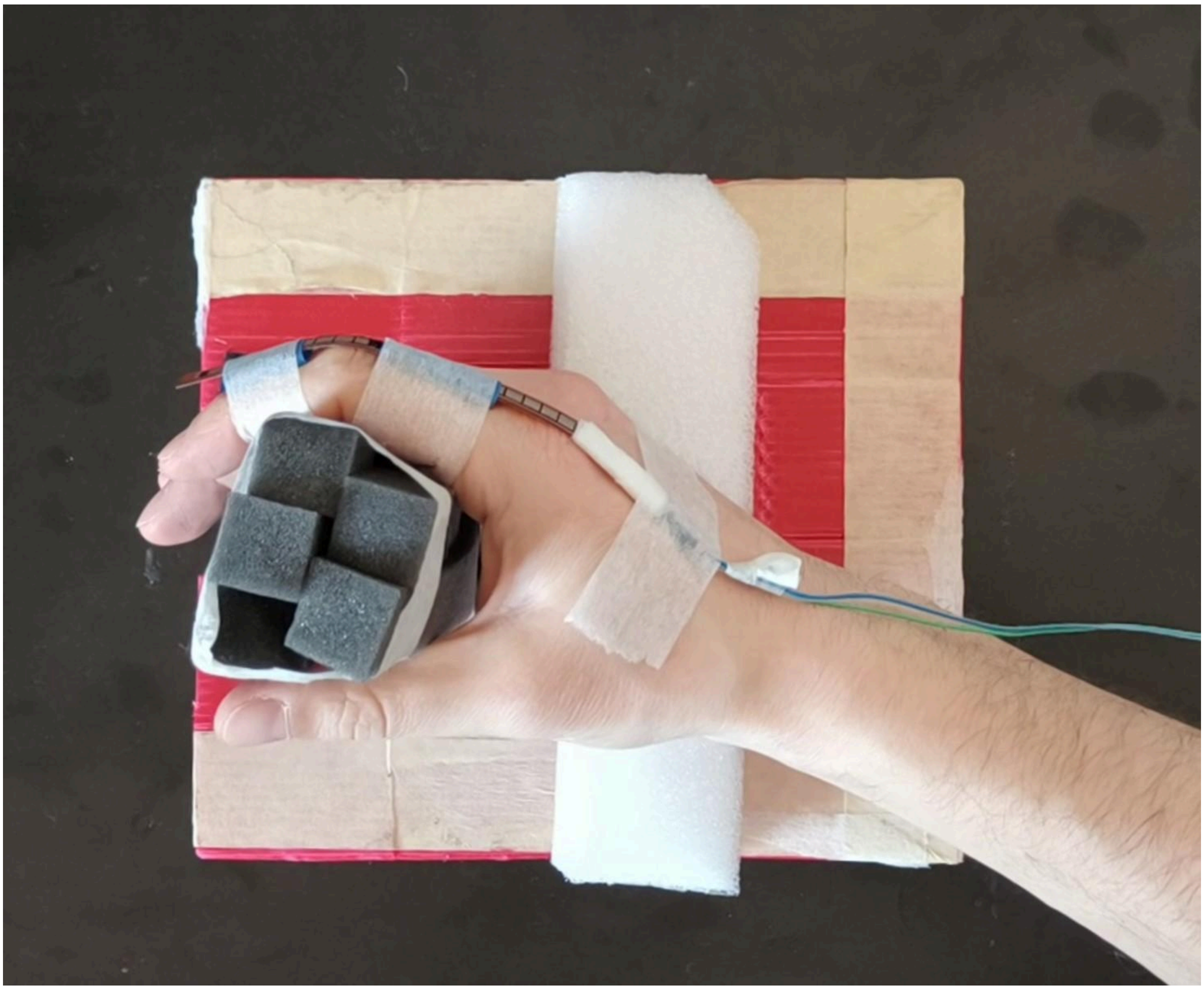

**Figure 1:** Example of a participant's hand starting position at the beginning of each trial. The foam rubber structure (dark grey) keeps the fingers in an intermediate position while waiting for the stimuli to appear on the screen. The hand lays on a rigid base (white), which allows the fingers to move freely during extension by creating space between the table and the ulnar side of the metacarpus. Additionally, if necessary, a soft foam rubber support has been placed beneath the forearm  to maintain alignment of the hand-forearm complex.

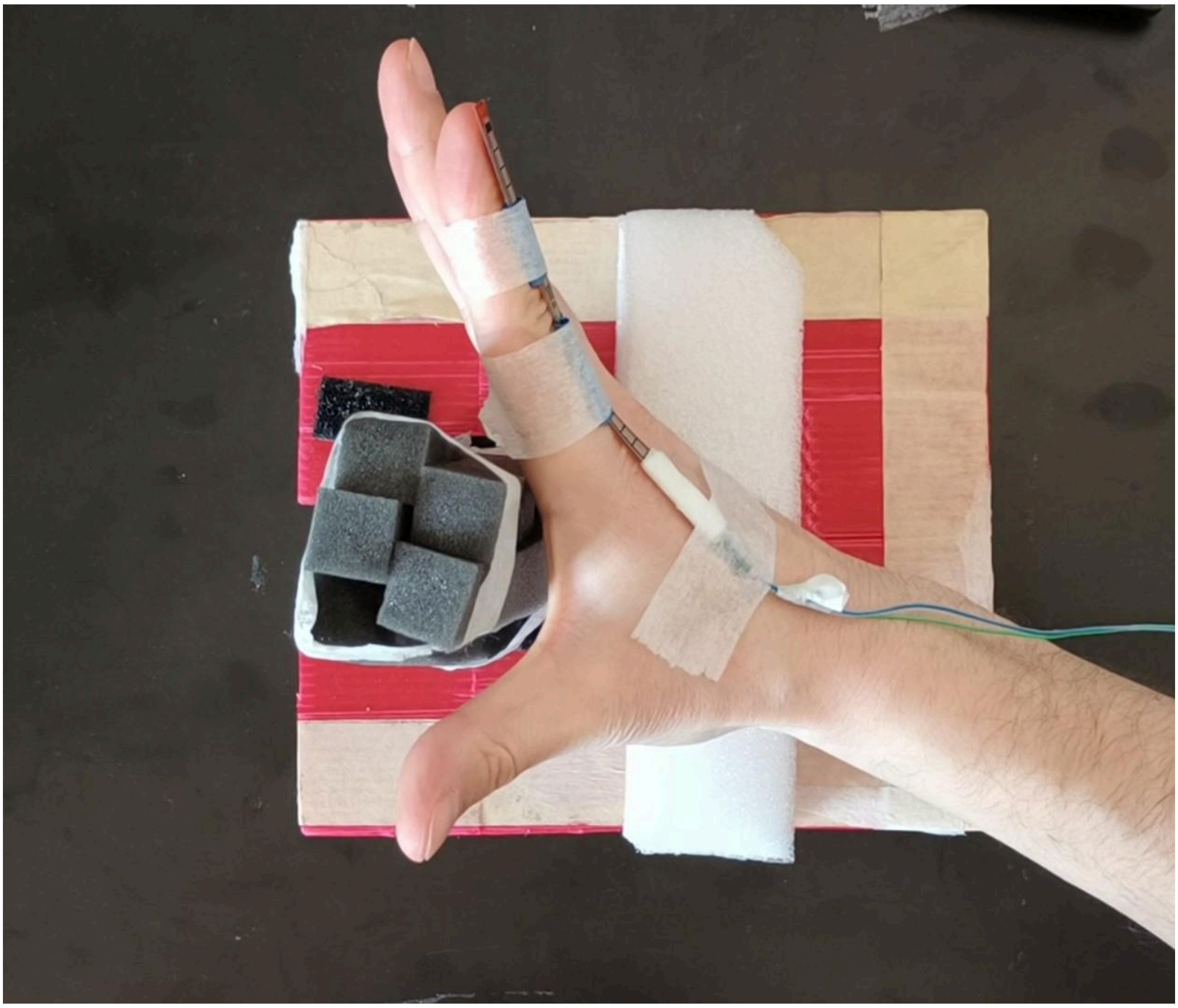

**Figure 2:** Example of fingers extension movement.

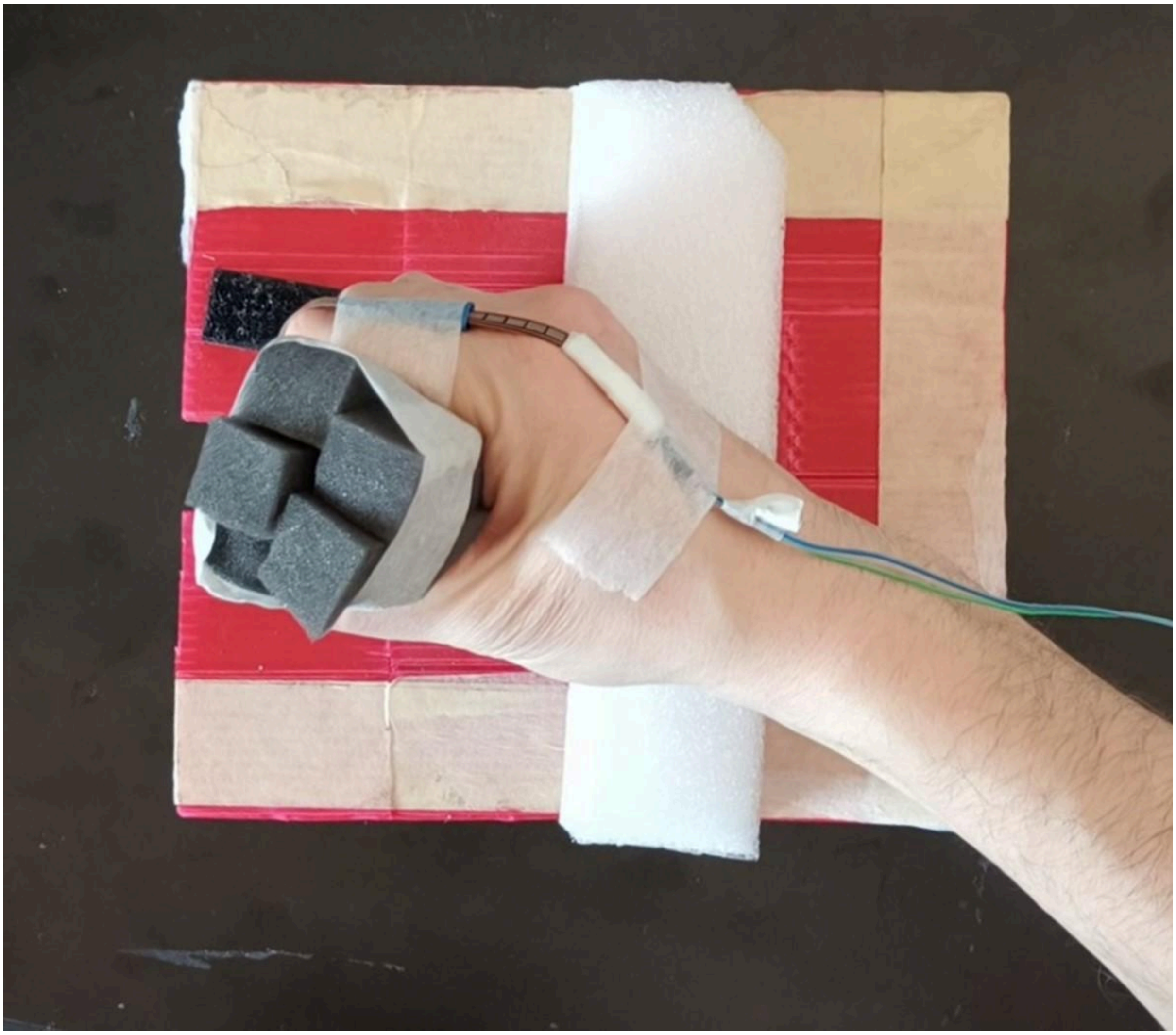

**Figure 3:** Example of fingers flexion movement.
